# KR158 spheres harboring slow-cycling cells recapitulate GBM features in an immunocompetent system

**DOI:** 10.1101/2024.01.26.577279

**Authors:** Avirup Chakraborty, Changlin Yang, Jesse L. Kresak, Aryeh Silver, Diana Feier, Guimei Tian, Michael Andrews, Olusegun O. Sobanjo, Ethan D. Hodge, Mia K. Engelbart, Jianping Huang, Jeffrey K. Harrison, Matthew R. Sarkisian, Duane A. Mitchell, Loic P. Deleyrolle

## Abstract

Glioblastoma (GBM) poses a significant challenge in clinical oncology due to its aggressive nature, heterogeneity, and resistance to therapies. Cancer stem cells (CSCs) play a critical role in GBM, particularly in treatment-resistance and tumor relapse, emphasizing the need to comprehend the mechanisms regulating these cells. Also, their multifaceted contributions to the tumor-microenvironment (TME) underline their significance, driven by their unique properties.

This study aimed to characterize glioblastoma stem cells (GSCs), specifically slow-cycling cells (SCCs), in an immunocompetent murine GBM model to explore their similarities with their human counterparts. Using the KR158 mouse model, we confirmed that SCCs isolated from this model exhibited key traits and functional properties akin to human SCCs. KR158 murine SCCs, expanded in the gliomasphere assay, demonstrated sphere forming ability, self-renewing capacity, positive tumorigenicity, enhanced stemness and resistance to chemotherapy.

Together, our findings validate the KR158 murine model as a framework to investigate GSCs and SCCs in GBM-pathology, and explore specifically the SCC-immune system communications, understand their role in disease progression, and evaluate the effect of therapeutic strategies targeting these specific connections.

## INTRODUCTION

GBM, the most common malignancy of the central nervous system, continues to present a significant challenge in oncology due to its aggressive nature and resistance to conventional therapies. The involvement of CSCs in GBM, especially in driving resistance to treatment and tumor relapse, makes it a priority to fully understand the mechanisms regulating this phenotype or cellular state. Extensive research has highlighted the multifaceted role of these cells, demonstrating their contribution to tumor initiation, therapeutic resistance, and recurrence. Their distinctive properties, notably their long-term self-renewal capacity and ability to generate a large number of progenies, underscore their influential role in driving the aggressive behavior of GBM.

Tumor heterogeneity is also a recognized major determinant of treatment failure and recurrence. Understanding the key mechanisms driving this heterogeneity is paramount to improve our understanding of the complex biology of GBM and design effective new therapies. CSCs are key to this heterogeneity and comprise diverse heterogeneous populations with multiple factors, both intrinsic and extrinsic, contributing to their phenotypic and functional diversity ^1-5^. Findings from our laboratory and others described CSCs in several malignancies ^6-17^, including GBM ^15-24^, where the existence of a subpopulation of CSCs, namely slow-cycling cells (SCCs), has gained attention due to their resistance to standard-of-care therapies and their role in GBM progression and recurrence. Despite their clinical relevance, effective strategies to eliminate these cells remain elusive. Hence, it is imperative to advance our understanding of their biology and identify vulnerabilities that can be exploited for therapeutic advancements.

The evolving comprehension of the role of the immune system within the GBM landscape has reshaped perspectives on the study of the TME, with recognition of immune responses, or lack thereof, emerging as a critical factor influencing clinical outcomes. This expanding understanding has prompted investigations into the dynamic interactions between cancer cells, especially CSCs and SCCs, and immune cells within the GBM microenvironment. Therefore, complexities of the TME, comprising diverse cell types and their intricate interactions, represent a focal point in unraveling GBM trajectory and treatments.

Developing and using pre-clinical GBM models adequately representing the TME, specifically mirroring its complexity, morpho-pathological characteristics, and immune landscape, is a prerequisite to integrate the dynamics of the different cellular components and their collaborative or antagonistic interactions. Deciphering the intricate relationships between tumor cells, e.g. SCCs, and the immune system and investigating the impact of tumor heterogeneity on the immune milieu has the potential to unveil pivotal mechanisms dictating disease progression and treatment responses.

One of the well-studied and used immunocompetent murine models of GBM to address these issues is KR158 ^25^. This model of glioma is driven by the mutation of two tumor-suppressor genes, NF1 and TP53 ^25^. Despite its relevance, this model has limitations in recapitulating human pathology, specifically in its tendency to mirror grade 3-like gliomas rather than grade 4-like tumors. Although orthotopic transplantation results in approximately 60% in malignant lesions displaying high-grade glioma features—such as multifocal lesions, varying cellularity, high mitotic index, and microscopic necrotic foci—it lacks extensive infiltration and the distinctive pseudopalisading around necrotic areas, thus falling short of classification as grade 4, or glioblastoma.

The gliomasphere assay was designed to enrich glioma stem cells capable of forming tumors that reproduce all key morphological traits of grade 4 glioma ^26,27^. In our study, we leveraged this assay to culture murine KR158 cells, aiming to isolate cellular populations proficient in recapitulating these distinctive characteristics. Our findings demonstrate that KR158 cells expanded under these conditions produce tumors resembling GBM when transplanted intracranially. These results support the efficacy of this assay in more closely recapitulating the human disease as compared to KR158 cells cultured in adherent monolayer serum-containing conditions. Our previous research established the existence of a metabolic diversity within the microenvironment of human GBM, which includes fast and slow cycling cells with distinct metabolic features ^1,5,16,19^. Moreover, SCCs isolated from human GBM tumors have demonstrated increased stemness, enhanced invasion, and drug resistance, pointing to their role in tumor progression and recurrence ^16,19,20^. The objective of this study was also to validate the slow/fast cycling cell paradigm in an immune competent model of glioma with the long-term goal of exploring the connections between the intratumor heterogeneity and the immune landscape in these tumors. This work showed that in the KR158 mouse model of glioma ^25^, SCCs share phenotypic and functional traits of their human counterparts ^16,19,20^. We found that KR158 SCCs are tumorigenic, generate tumors displaying key features of GBM, exhibit a greater stemness phenotype and specific metabolic signature, and enhanced resistance to chemotherapy. Together, our results validate the potential use of this model for a comprehensive exploration of the intricate connections between CSCs, especially SCCs, the immune system, and the TME in GBM. This exploration will emphasize their profound implications for disease understanding and the development of innovative therapeutic strategies.

## MATERIALS AND METHODS

### Murine glioma cell line

Murine KR158 (wildtype or expressing luciferase [Kluc]) glioma cell line was kindly provided by Dr. Tyler Jacks, Massachusetts Institute of Technology, Cambridge, MA, USA ^25^. These cells were cultured both in serum-free gliomasphere assay conditions as floating spheres and in standard adherent monolayer and serum containing conditions as described below.

### Gliomasphere assay

Under serum-free conditions, the cells were cultured at 37□°C in the presence of 5% CO_2_ in NeuroCult NS-A Proliferation solution with 10% proliferation supplement (STEMCELL Technologies, Vancouver, BC, Canada; Cat# 05750 and #05753) supplemented with 20ng/mL mouse EGF (R&D Systems, Minneapolis, MN, USA, Cat#2028-EG),10 ng/mL human FGF2 (Catalog Number: 233-FB/CF, R&D Systems, Minneapolis, MN, USA), which stimulates the proliferation of murine cells ^28^. Human and murine FGF receptors are highly conserved and show a 95% homology ^29^. We also included 10ng/mL of Heparin (Sigma-Aldrich, St. Louis, MO, USA, Cat#H3149) as it regulates FGF2 activity, enhances specificity and affinity towards the FGF2 receptor, thereby modulating the transduction cascade and stimulating cell proliferation ^30-32^. Cultures were maintained in 1% Antibiotic-Antimycotic (Life Technologies, Carlsbad California, USA, Cat# 15240062). Upon reaching a diameter of about 150 μm in diameter, gliomaspheres were enzymatically digested using Accutase (StemCell Technologies, Vancouver, BC, Canada Cat#07920) for 15 min at 37°C. Subsequently, the cells were washed, counted, and replated in a fresh serum-free complete medium for further expansion.

### Classical adherent and serum-containing cell cultures

The cells were cultured at 37□°C in the presence of 5% CO_2_ in DMEM (Gibco, New York, USA Cat#11965092) and supplemented with 10% FBS (Avantor, Pennsylvania, USA Cat#89510186) and 1% Antibiotic-Antimycotic (Life Technologies, Waltham, MA, USA, Cat# 15240062). Upon reaching 90% confluency, the cells were enzymatically digested using Accutase (StemCell Technologies, Vancouver, BC, Canada Cat#07920) for 15 min at 37°C, subsequently were washed, and replated in fresh complete medium.

### Scratch-Wound Assay and Time⍰lapse imaging

Cells that were expanded in the gliomasphere assay for several passages were plated in a 35 mm time-lapse imaging dish. Attachment of the cells for this assay was stimulated by the addition of 10% FBS. Imaging was performed on an inverted Zeiss Axio-Observer D1 microscope using a Zeiss. Imaging dishes were secured onto a stage-top incubation system and maintained in a humid chamber at 37 °C and 5% CO_2_ using a Tokai Hit System. Image acquisition and processing were performed using the Zeiss ZEN software. Images were acquired every 10 minutes and movies were exported at 5 frames per second.

### Intracranial implant, tumor growth monitoring and survival analysis

To compare the tumor generating capabilities of the i) total unsorted Kluc cells grown as gliomaspheres against those grown as adherent cells and ii) total unsorted Kluc, FCCs and SCCs grown in the gliomasphere assay, the tumor cells were intracranially implanted in immune competent 7–15-week-old C57BL/6 mice, following NIH and institutional (IACUC) guidelines and regulations for animal care and handling. The mice colonies were maintained at the University of Florida’s animal facility. These mice were intracranially implanted with 2μl of cell suspension containing 10,000 live Kluc cells using a sterile 5 ml Hamilton syringe fitted with a 25-gauge needle into the striatum using a stereotactic apparatus. The injection coordinates were 2.0-mm lateral to the bregma at a depth of 3.0 mm below the dura mater as previously described ^16,19,33^. Longitudinal monitoring of the tumor volume was performed using an IVIS Spectrum imaging system (Xenogen, Alameda, CA) measuring bioluminescence related to luciferase activity. The animals were also monitored for any neurological signs affecting their quality of life. When symptoms including ataxia, lethargy, seizures or paralysis were observed, the mice were sacrificed.

### Isolation of fast and slow cycling cells

SCCs and FCCs cell population grown as glioma spheres were identified and isolated based on their proliferation rate which was accessed based on their ability to retain carboxyfluorescein succinimidyl ester— CFSE [Cat#c1157] or Cell Trace Violet-CTV [Cat#c34571], (Invitrogen, Waltham, MA, USA), as described previously ^16,17,19^ by studying the CFSE/CTV fluorescence intensity decay rate overtime as measured by flow cytometry. 7 days post labeling, these cells were grouped as CFSE/CTV^High^-top 10% (referred to as SCCs) and CFSE/ CTV^Low^-bottom 10% (referred to as FCCs). This gating strategy allowed us to isolate the functional and phenotypic extremes with similar-sized populations, homogenizing for sorting time and addressing potential issues related to fluorescence-activated cell sorting (FACS)-induced metabolic stress. The utilization of these extreme fractions from the proliferation spectrum ensured a clear and distinct separation of FCCs and SCCs based on cell cycle kinetics. All experiments were promptly conducted following the FACS of SCC and FCC populations.

### Brain tumor tissue processing

Upon reaching the defined humane endpoint or upon reaching the designated study time point, the mice were euthanized, and the tumor tissues were resected and frozen in OCT or preserved in paraffin. The preserved tissues were then sectioned into 5-8μm slices and placed on slides. *In situ* tumor formation with classical morphological features of GBM including infiltration, nuclear pleomorphism with mitotic figures, and pseudopalisading necrosis was confirmed using hematoxylin and eosin staining of the paraffin-embedded sections. Further, we determined *in vivo* the invasiveness of the luciferase-expressing tumor cells by staining the frozen OCT-embedded sections with anti-Firefly Luciferase antibody (1:500 Abcam, Cambridge, UK Cat#21176 respectively) and a goat anti-rabbit AF488 conjugated secondary antibody (1:1000; Cat#11008 Invitrogen, Waltham, MA, USA).

### Live/dead assay

Propidium Iodide (PI) incorporation assay using flow cytometry: To determine the impact of TMZ on the viability of the SCCs, FCCs, and the total unsorted tumor cells, we performed an *in vitro* PI incorporation assay. The cells were stained with CTV and expanded for 7 days before being exposed to a dosage of 400μM of TMZ for 48 hrs. Their viability was assessed using PI staining (Thermo Scientific; Waltham, MA, USA). Cell staining was performed with the method as recommended by the manufacturer’s manual. The incorporated PI was quantified using flow cytometry (BD LSRII; BD Biosciences Franklin Lakes, NJ, USA). The cells were gated with respect to the intensity of CTV and identified as SCCs and FCCs as described above. PI positive cells were indicative of dead cells. The value of PI positive cells represents the mean of three independent experiments.

### CyQUANT Direct Cell Proliferation Assay

The cytotoxic effects of TMZ at a dosage of 2mM was further assessed by using the fluorescence-based CyQUANT Cell Proliferation assay (Thermo Fisher Scientific Waltham, MA, USA Cat#C7026). The assay measures proliferation and membrane integrity, another measure of cell health. Sorted SCCs, FCCs and unsorted tumor cells were plated at 80,000 cells per well in 96-well plates and exposed to 2mM TMZ. After 48 hrs, CyQUANT binding dye was added to each well and incubated for 30 min at 37°C before being quantified using BiotekTM Cytation™ 3 Cell Imaging Multi-Mode Reader. The value of relative cell proliferation represents the mean of three independent experiments.

### Sphere forming frequency assay

To determine the self-renewal nature of the cancer stem cell fraction, we performed the sphere forming frequency assay of the sorted SCCs, FCCs and the unsorted tumor cells when exposed to 1mM TMZ. To achieve this, we added a single cell suspension of 1000 cells to each well of the 96 well plate containing the TMZ dosage and incubated them for 4 days. The cells were then fixed, permeabilized and the nuclei were stained using a solution with a final concentration of 2% PFA (Thermo Scientific; Waltham, MA, USA), 0.01% Triton™ (Roche; Basel, Switzerland) in PBS, and 0.1% DAPI (Thermo Scientific; Waltham, MA, USA). The Gen5 Image software on the Cytation™ 3 Cell Imaging Multi-Mode Reader was utilized to quantify spheres that exhibited a circularity exceeding 0.15 and measured between 50 to 500 μm in diameter.

### Bulk RNA-Seq sample preparation and analysis

RNA extraction, RNA-seq library generation, and sequencing were conducted following previously established protocol ^18^. In summary, total RNA was extracted from cells cultured under adherent or serum-free gliomasphere assay conditions and from brain tumor tissue samples using Qiagen kits (Toronto, ON, Canada). Prior to library construction, rigorous quality control steps were implemented to ensure RNA purity and integrity. This involved initial assessment of RNA purity using Nanodrop2000 (Thermo Scientific; Waltham, MA, USA), agarose gel electrophoresis for RNA integrity and potential contamination, and reconfirmation of RNA integrity using the Agilent 2100 Bioanalyzer (Santa Clara, California, USA). Subsequently, mRNA was purified and randomly fragmented to initiate cDNA synthesis. For library construction, cDNA fragments of 150-200bp length were purified to a concentration of 1.5ng/μl using AMPure XP beads (Beckman Coulter, Beverly, USA). Assessment of library effective concentration and RNA quality were performed using Agilent 2100 Bioanalyzer and Qubit2.0 (Thermo Scientific; Waltham, MA, USA). Finally, the libraries underwent sequencing on the HiSeq platform (Illumina, San Diago, CA, USA) following the manufacturer’s protocols. The assessment of gene signature enrichment was carried out using GSEA (http://www.broadinstitute.org/gsea/index.jsp). The stem cell signatures utilized in this study were derived from Wong et. al. ^34^ and Harris et. al. ^35^. The SCC signature was previously described by Hoang-Minh *et al*. and Yang *et. al*. ^19,20^. The GO terms used for the cell migration, cell motility, response to lipid, and lipid catabolism signatures were: GO:0016477, GO:0048870, GO:0071396 and GO:0016042, respectively. The KEGG pathway mmu04512 was used as ECM receptor signature. Nominal p value < 0.05 and FDR<0.25 were applied to detect gene set enrichment across different groups. Multivariate principal component analysis (PCA) was utilized to differentiate between tumors derived from SCC or FCC and to compare them with control brain tissues, examining the transcriptomic diversity by using FactoMineR. Differential expressed genes (DEGs) were extracted by using R limma (logFC > 1.5 or < −1.5 and BH adjusted p value < 0.01) between KR158 derived tumors and GL261 derived tumors and their hierarchical clustering analysis was performed by using R pheatmap.

Data presented in this paper is available on BioProject database (PRJNA1068559).

### Single-cell RNA sequencing and quality control

CD45 negative cells from KR158 intracranial tumors were isolated and processed for scRNA sequencing as described by Trivedi *et. al*. (*doi: 10.1186/s13073-024-01281-z*). In brief, CD45 negative cells were obtained as the flow-through fraction after isolating CD45 positive cells using bead-based selection. A cDNA sequencing library was then prepared using the Chromium Next GEM Single Cell 3L Reagent Kits v3.1 (Dual Index, 10x Genomics) and sequenced on an Illumina Novaseq 6000. For downstream analysis, Trivedi *et al* only used samples with a Cell Multiplexing Oligo assignment probability above 70% and a multiplet rate below 50% and retained genes that were expressed in at least three cells. Cells with more than 250 genes, 500 UMIs, a Complexity above 0.8 (log10 gene count/log10 UMI count), and less than 5% mitochondrial genes were selected using Seurat version 4.045.

### Single-cell RNA-seq data analysis

We utilized SingleR, integrated with the Immgen dataset, to identify cell types (PMID: 30643263) from the scRNAseq dataset obtained. Seurat 4.0 was employed to identify Differentially Expressed Genes (DEGs) within cell clusters for subsequent cell type classification. 1,000 CD45 negative cells were included in the analysis. We visualized cell populations using the UMAP algorithm via Seurat 4.0. SCCs and FCC were defined using the genes signature and score as previously described ^19,20^. Stemness geneset ^34^ and Cell motility (GO:0048870) scores in SCCs were determined by using the Escape package. UMAP projections of scores were generated using the FeaturePlot function in Seurat 4.0.

### Statistical analyses

The results are expressed as mean values ± SEM, with statistical analyses performed using GraphPad Prism 6.0 (GraphPad Software). Statistical tests are indicated in the text. Group comparisons involved either a one-way ANOVA or Student’s t-test with 95% confidence intervals. ANOVA-significant groups underwent Tukey’s post hoc analysis. Overall survival was analyzed and compared using log-rank analyses. The flow cytometry results were analyzed using FlowJo™ Software (BD Life Sciences; Franklin Lakes, New Jersey, USA).

## RESULTS

### Transcriptomic difference between KR158 and GL261, with KR158 overexpressing GBM-like signatures

GL261 is one of the most predominantly used preclinical models for GBM ^36^. However, allograft tumors generated by this model exhibit features including low clonotypic diversity and high antigenicity, contrasting with human GBM ^37^. The murine model KR158 is another commonly used immunocompetent model of glioma, which exhibits several properties that closely mirror those observed in human GBM ^38^. In our effort to assess the capacity of both models to replicate characteristics resembling GBM, we first conducted a multivariate principal component analysis (PCA) on tumors originating from KR158 and GL261 cells. Our analysis revealed transcriptomic differences between these distinct tumor cell populations **(*Fig. 1A*)**. The heatmap representation of hierarchical clustering analysis on the DEGs further underscored the transcriptomic distinctions among these tumor models (***Fig.1B***). Given the significance of stemness and migration in human GBM ^39-44^, we compared the expression levels of various genesets associated with these properties in tumors originating from GL261 and KR158. Analysis using Gene Set Enrichment Analysis (GSEA) on bulk RNA sequencing data indicated elevated expression of stemness and cell migration-related genes in tumors derived from KR158 compared to those from GL261 **(*Fig. 1C-F, Tables 1-5*)**. Given our goal to investigate a specific subset of cells, namely SCCs, in an immune-competent model of GBM, it was critical to examine the transcriptomic profile of these models in relation to genes previously associated with such cells or cellular states in human GBM ^19,20^. Interestingly, our findings revealed an up-regulation of the SCC gene signature in tumors derived from KR158 as compared to those from GL261 (***Fig. 1G***). This observation suggests that the KR158 model may be more relevant when studying this population of GBM SCCs.

**Figure 1.**
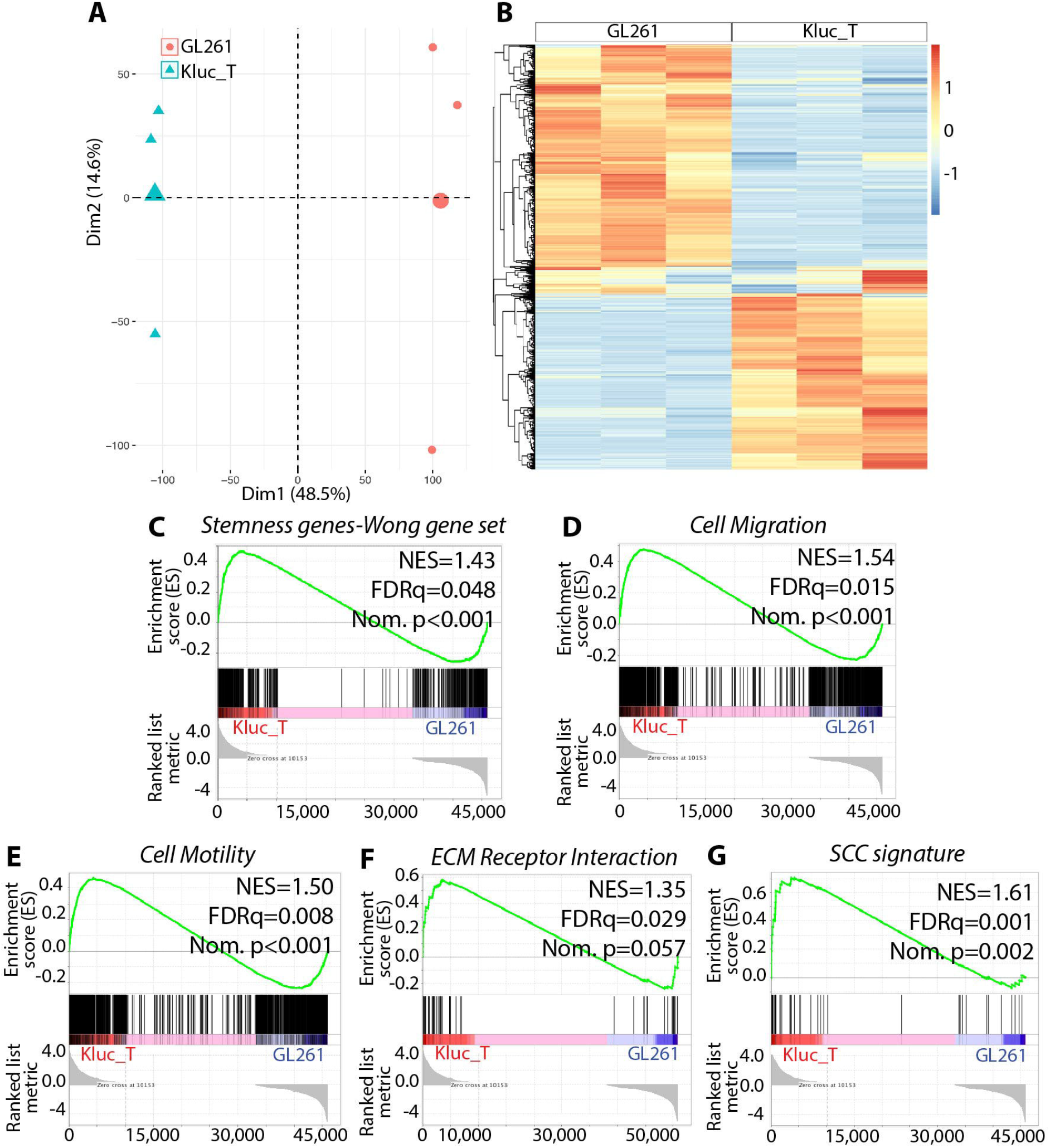
Differential transcriptomic profiles of kKR158 and GL261, highlighting upregulation of genes associated with GBM-like features in KR158. **A)** PCA score plot from bulk RNA sequencing performed on tumors derived from GL261 and KR158 brain tissue (n=3), showed that these tumor models exhibit different transcriptomic profiles. **B)** Heatmap showing DEGs of tumors derived from GL261 and KR158 brain tissue (n=3). Red and blue indicate relative over-or under-expression of genes, respectively. **C-G)** GSEA between tumors derived from KR158 and GL261 brain tissue for the following genesets (n=3 per group); (**C**) stemness (signature from Wong *et al* ^*34*^), (**D**) cell migration (GO:0016477), (**E)** cell motility (GO:0048870), (**F)** ECM receptor interaction (KEGG mmu04512), (**G)** SCC gene signatures ^19,20^; FDR, false discovery rate; NES, normalized enrichment score; Nom., nominal.

### scRNA sequencing revealed cellular diversity in KR158 tumors, exhibiting similarities with the heterogeneity observed in hGBM

Utilizing a single-cell RNA (scRNA) sequencing dataset obtained from KR158 intracranial tumors (*doi: 10.1186/s13073-024-01281-z*), we analyzed gene expression in a thousand excised cells. These cells were classified into eight distinct cellular clusters, each representing a major cell type, including tumor cells, microglia, endothelial cells, oligodendrocytes, astrocytes, fibroblasts, neurons, and epithelial cells (***Fig. 2A***). This result illustrates the cellular heterogeneity within the tumor microenvironment of this model, a characteristic also identified in human disease. The tumor cell group was subdivided into two distinct subpopulations—namely, the SCC and FCC cellular clusters—distinguished by the expression of specific gene signatures as previously described in hGBM ^20^. Notably, the SCC population displayed significant heterogeneity, evident in its widespread distribution on the UMAP plot, contrasting with the more compact cluster formed by the FCC cells (***Fig. 2A***). Additionally, in alignment with our observations in hGBM, the SCCs demonstrate elevated expression of genes associated with stemness and cellular motility, which are key features of hGBM (***Fig. 2B-C***) ^45,46^. Collectively, these results support the significance of the KR158 model for unraveling the tumor microenvironment and exploring the characteristics of SCCs.

**Figure 2.**
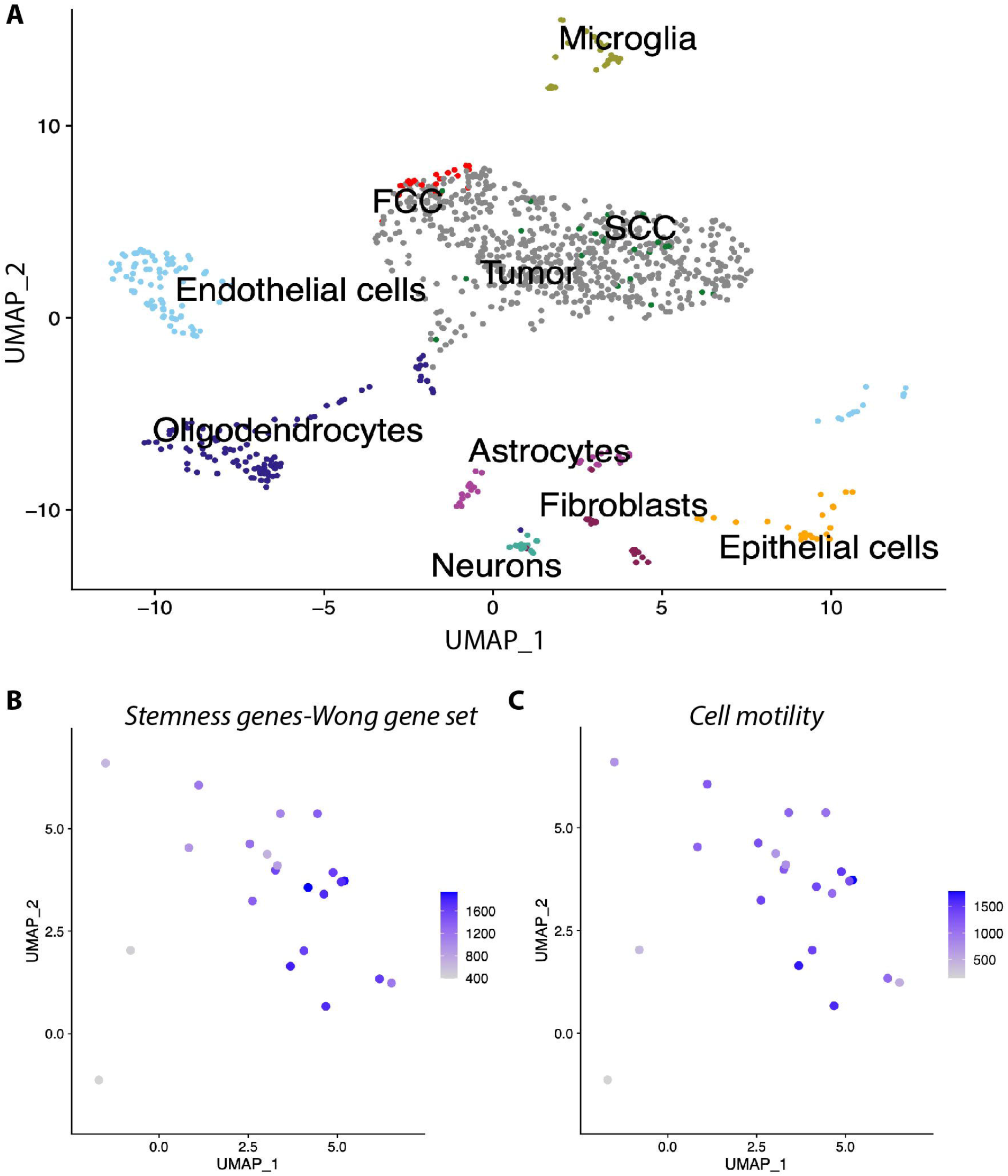
scRNA sequencing revealing cellular diversity in KR158 tumors. **A)** UMAP visualization of pooled scRNA-seq data of 1000 CD45 negative cells of the tumor microenvironment from tumors derived from KR158 brain tissue. We identified 8 clusters, including tumor cells, microglia, endothelial cells, oligodendrocytes, astrocytes, fibroblasts, neurons, and epithelial cells. Tumor cell fraction contains subpopulation of SCCs (green) (22 cells) and FCCs (red) (22 events). **B-C)** Stemness geneset ^34^ (**B**) and Cell motility (GO:0048870) scores (**C**) of each single cell were determined by using the Escape package. Scores were protected to UMAP by using Seurat 4.0 FeaturePlot function.

### KR158 cells expanded using the gliomasphere assay generate glioblastoma-like tumors

With the goal to support the cancer stem cell phenotype in an immunocompetent model of GBM, KR158 cells were cultured using the serum-free gliomasphere assay ^25,27^. Under these conditions, KR158 cells formed spheres (***Fig. 3A***), as opposed to an adherent monolayer growth pattern when cultured in standard conditions containing serum (***Fig. 3B***). The scratch-wound assay demonstrated that cells expanded under the defined gliomasphere conditions maintain their migration capacity, a critical characteristic of GBM cells (***Fig. 3C-D, Supp. video 1***). Importantly, orthotopic transplantation of a single cell suspension of luciferase-expressing KR158 cells expanded in the gliomasphere assay led to the development of brain tumors (***Fig. 3E***). These tumors exhibited hallmark features of grade 4 glioma, such as extensive invasion with notable subpial spread (***Fig. 3F-G***). Cellular and nuclear pleomorphism with giant cells were also observed along with abundant mitosis and perivascular aggregates (***Fig. 3H-h***). The presence of pseudopalisading necrotic regions characterized by a hyper cellular nuclei rearrangement surrounding irregular foci of tumor necrosis containing pyknotic nuclei were also observed (***Fig. 3I***). Notably, tumors originating from cells grown in monolayer serum-containing conditions also exhibited evidence of necrosis, albeit with less defined palisades and a sparser cellular arrangement surrounding the necrotic foci (***Fig. 3J***). Furthermore, tumors derived from serum-containing cultures exhibited a less extensive infiltrative phenotype, displaying a more circumscribed growth pattern (***Fig. 3K***). These findings highlight the tumorigenic nature of KR158-derived gliomaspheres, generating tumors that exhibit characteristics more akin to grade 4 tumors. This distinction is evident in the presence of high infiltration and well-defined pseudopalisading necrosis, unlike the grade 3-like malignancies lacking such features when cultured under adherent serum-containing conditions.

**Figure 3.**
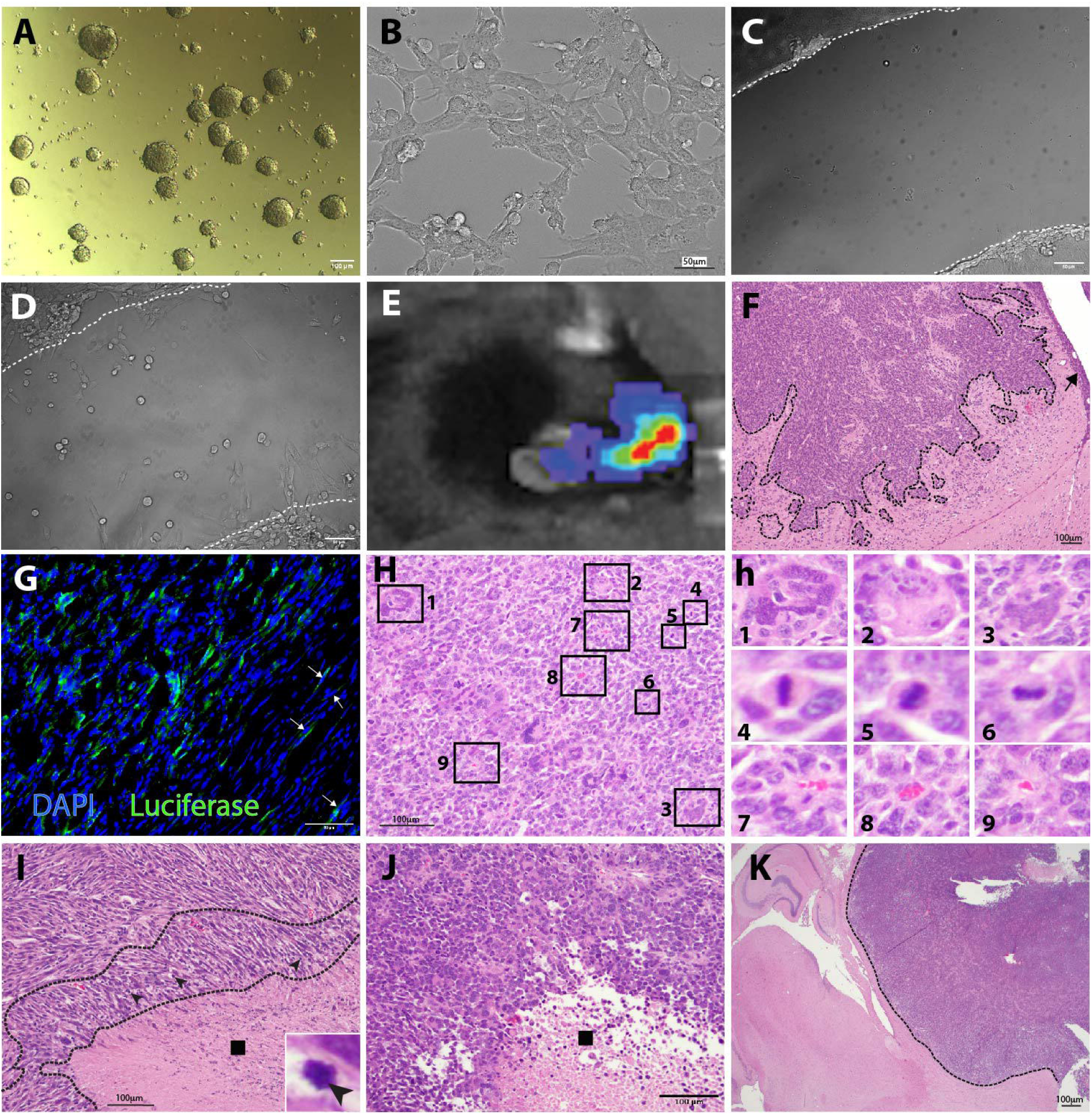
KR158 murine glioma cells grown in the glioma sphere assay demonstrate tumor formation characterized by hallmarks of high-grade glioma. **A)** Brightfield image of Luciferase-expressing KR158 cells (KLuc) murine glioma cells grow as spheres when cultured in serum-free medium supplemented with epidermal growth factor (EGF) and basic human fibroblast growth factor (hFGF). **B)** Brightfield image of KLuc cells cultured in adherent and serum-containing conditions. **C-D)** Scratch-wound assay; brightfield images acquired at time 0 (**C**) and 23h (**D**) show the migratory behavior of the cells that were expanded in the gliomasphere assay. **E)** Luciferase-based in vivo imaging using IVIS, of tumor-bearing C57BL6 mice when intracranially implanted with KLuc cells. F) Hematoxylin and eosin staining (H&E) show that intracranial implantation with KLuc cells cultured in serum-free conditions in C57BL6 mice demonstrate tumorigenicity with ability to generate tumors exhibiting GBM characteristics including infiltration. Tumor leading edge denoted by the dotted line, → indicates subpial spreading. **G)** Luciferase labeling (green) further confirms invasive properties of KLuc into the host brain parenchyma. White arrowheads indicate luciferase^+^ cells that have migrated away from the tumor core, infiltrating the surrounding brain parenchyma. Nuclei are labeled with DAPI (blue). **H**) H&E staining of tumors developed from cells cultured in the gliomasphere assay depicts the presence of giant cells, identified as # 1-3, mitotic figures (#4-6), and clustering in perivascular regions (#7-9). Panel (**h**) represents higher magnification insets indicated by the rectangles in panel H. **I**) The presence of pseudopalisading necrosis further validates the formation of high-grade glioma like disease from KLuc cells expanded in gliomasphere serum-free medium. L indicates necrotic region, dotted line indicates pseudopalisade, l1 indicates pyknotic nuclei. **J-K**) H&E labeling of brain sections of animals implanted with KLuc cells cultured in serum containing conditions. Images presenting pseudopalisading necrosis (**J**) with more limited infiltration (**K**) compared to tumors derived from cells cultured in serum-free conditions. Tumor leading edge denoted by the dotted line. Shown are representative images from 5 mice per group.

### The gliomasphere assay enhances stemness gene signature in KR158 cells

Upon confirming through histopathological evidence that KR158-derived gliomasphere tumors align better with grade 4 tumor characteristics, we compared the transcriptomic profiles of KR158 cells grown in gliomasphere and adherent conditions. Utilizing multivarient PCA from bulk RNA sequencing, we identified notable transcriptomic diversities between these two cell populations (***Fig.4A***). This difference was further characterized by hierarchical clustering analysis of DEGs, visualized in a heatmap (***Fig. 4B***). GSEA indicated an increased stemness in cells grown in the gliomasphere assay, evidenced by an enriched stem cell gene signature in the tumors generated from these cells when compared to those grown under monolayer serum-containing conditions (***Fig. 4C-D, Tables 6-7***) ^34,35^. Utilizing the STRING platform (https://string-db.org/) for network analysis, it was observed that the genes showing significant up-regulation in cells cultured via the gliomasphere assay, as indicated in Tables 6-7, formed a network associated with the regulation of nervous system development (***Fig. 4E, Table 8***). Examples of genes present in this network associate with stem cells and the regulation of their self-renewal and differentiation, such as Mycn ^47-51^, Sox6 ^52^, Ncam2 ^53^, Fgf1 ^54-56^, Kit ^57^, and Larp6 ^58-60^.

**Figure 4.**
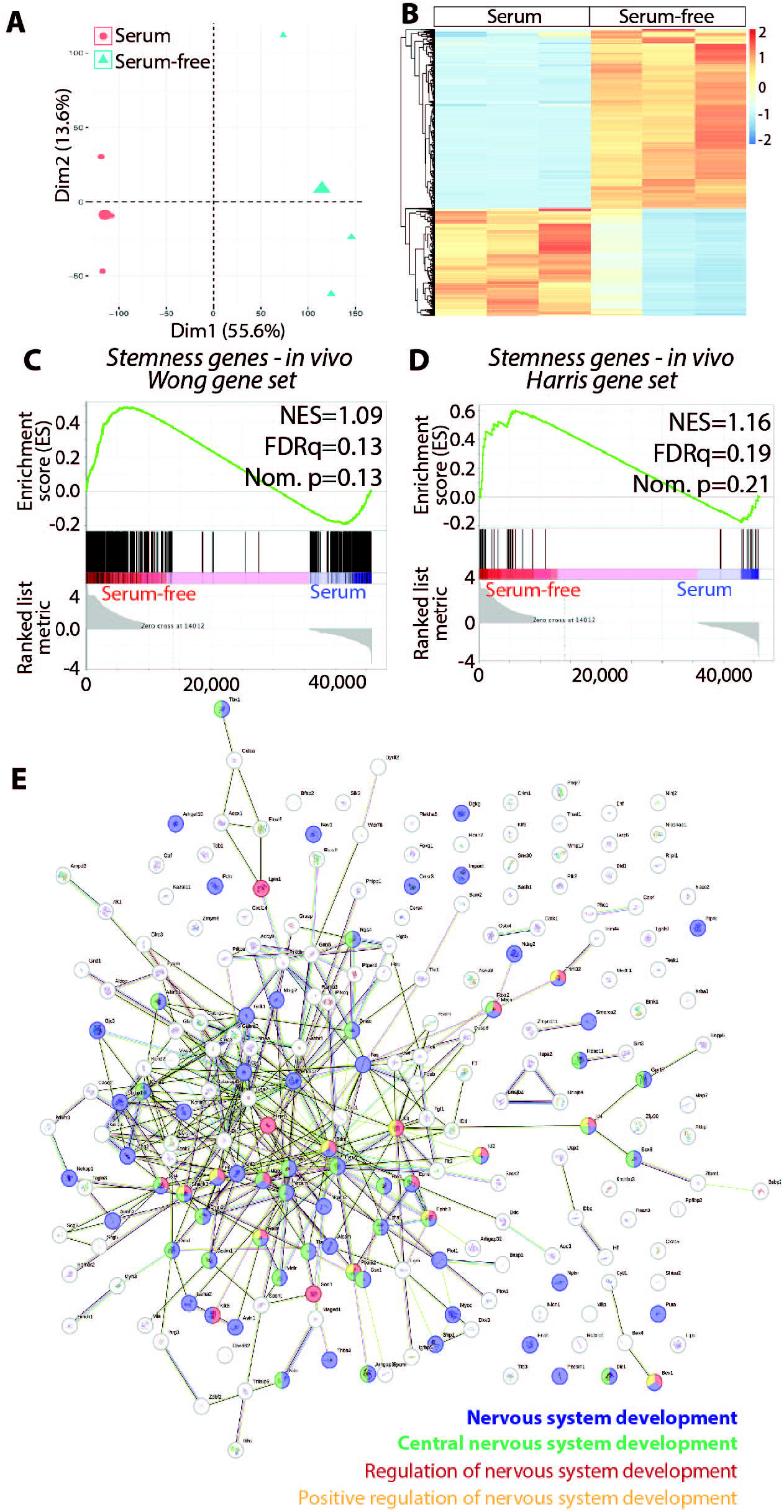
KR158 murine glioma cells grown in serum-free medium are enriched in Stemness genes. **A)** PCA score plot from bulk RNA sequencing performed on brain tumor tissue generated by KR158 cells cultured adherent in serum or serum-free gliomasphere assay (n=3), showed that these tumor models exhibit different transcriptomic profiles. **B)** Heatmap showing DEGs of tumors formed by KR158 cells cultured in serum or serum-free gliomasphere assay (n=3). Red and blue indicate relative over-or under-expression of genes, respectively. **C-D)** GSEA of RNA sequencing data from *in vivo* tumors (n=3 per group) derived from KR158 cells cultured in serum-free or serum-containing conditions shows an enrichment of stemness gene signature in tumors generated by cells cultured in serum-free conditions compared to cells expanded in serum-containing conditions using gene signature from **C**: Wong *et. al*. ^*34*^ and **D**: Harris Brain Cancer Progenitors gene set ^35^, systematic name M1694; FDR, false discovery rate; NES, normalized enrichment score; Nom., nominal. **E**) STRING (Search Tool for the Retrieval of Interacting Genes/Proteins) database analysis identified computational predictions defining functional associations between the genes upregulated in the KR158 cells expanded in the gliomasphere assay and mechanisms regulating the development of the nervous system.

### Evidence of SCCs in KR158 gliomasphere cultures showing similarities to SCCs in hGBM

Considering their ability to recapitulate the histological features of human GBM and their heightened stemness, KR158 cells grown in the gliomasphere assay were then used to identify and characterize SCCs in comparison to FCCs. KR158 SCCs were defined as CellTrace dye-retaining cells (***Fig. 5A-B***), as per previously established method ^16,17,19,20^. Bulk RNA sequencing showed differential clustering between the different tumor types through PCA (***Fig. 5C***), which is also visualized through heatmap representing the hierarchical clustering analysis of differentially expressed genes (***Fig. 5D***). Consistent with our findings in human GBM ^19^, GSEA from RNA sequencing showed that stemness genes were expressed at higher levels in murine glioma SCCs compared to FCCs when cultured in the gliomasphere assay (***Fig. 5E-F, Table 9-10***) along with genes corresponding to SCC signature (***Fig. 5G, Table 11***), cell migration (***Fig. 5H, Table 12***), cell motility (***Fig. 5I, Table 13***), and ECM receptor interaction (***Fig. 5J, Table 14***). Additionally, we found that lipid catabolic processes and response to lipid related genes were also up-regulated in SCCs, supporting a distinct metabolic profile similar to what we previously described in human GBM cells (***Fig. 5K-L, Tables 15-16)*** ^19^.

**Figure 5.**
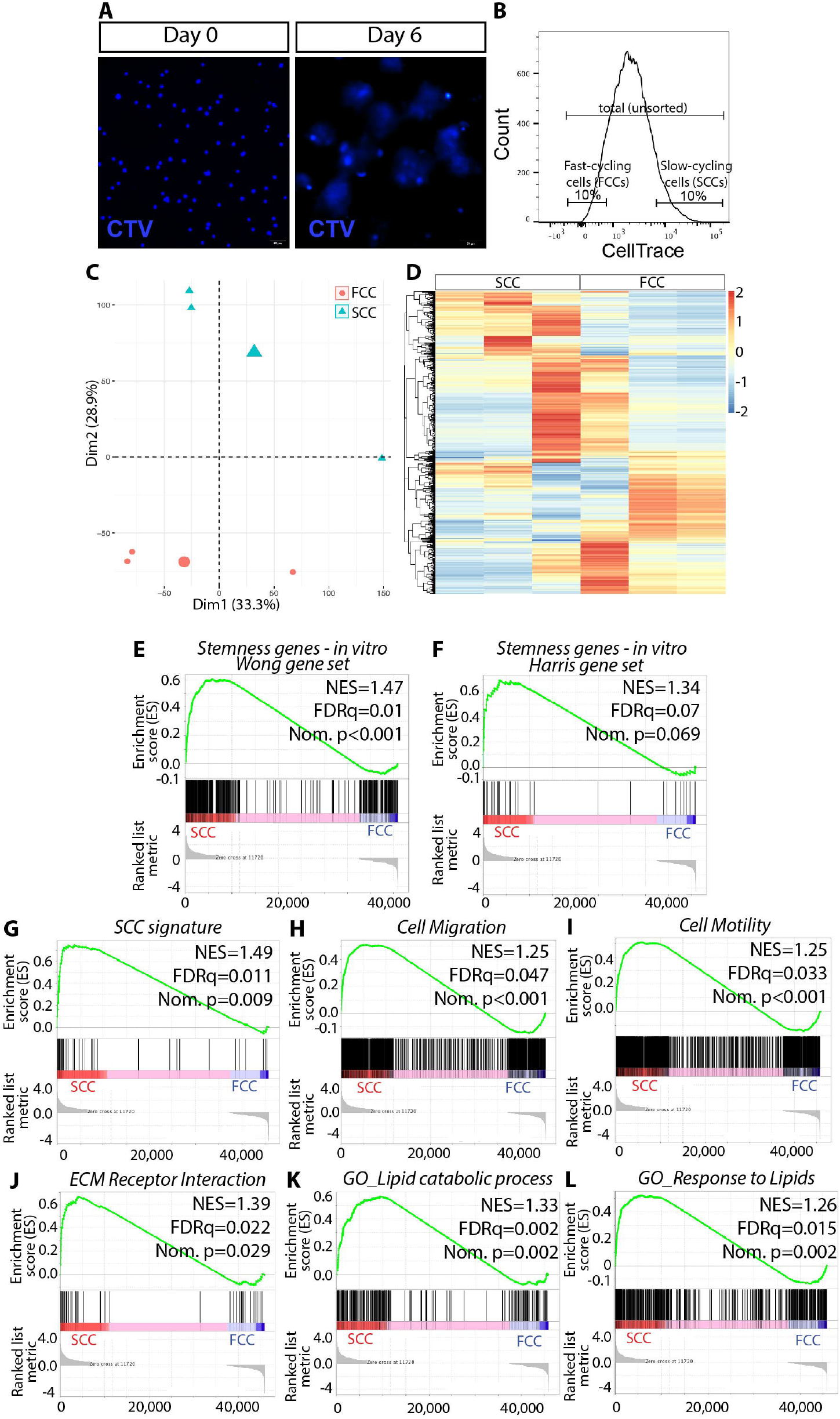
SCC KR158 cells grown in serum-free conditions are enriched in Stemness genes and lipid catabolic gene signatures over FCC cells. **A)** Cell trace labeling of KR158 cells grown in the gliomasphere assay at Day 0 and Day 6. **B)** Fluorescent-activated cell sorting of KLuc cells were performed on Day 6; top10% of the cells with highest CTV retention were sorted and defined as SCCs; and the bottom 10% of cells with least CTV retention as FCCs. **C)** PCA score plot from bulk RNA sequencing performed on tumors derived from SCC and FCC brain tissue (n=3), showed that these tumor models exhibit different transcriptomic profiles. **D)** Heatmap showing DEGs of tumors derived from SCC and FCC brain tissue (n=3). Red and blue indicate relative over-or under-expression of genes, respectively. GSEA of *in vitro* RNA seq data sets between gliomasphere serum-free cultured SCC (n=3) and FCC (n=3): (**E**) stemness geneset ^34^, (**F**) stemness geneset (M1694) ^35^; (**G**) SCC gene signature ^19,20^, (**H**) cell migration (GO:0016477), (**I**) cell motility (GO:0048870), (**J**) ECM receptor interaction (KEGG mmu04512), (**K**) lipid catabolic process (GO:0016042), (**L**) response to lipid (GO:0071396). FDR, false discovery rate; NES, normalized enrichment score; Nom., nominal.

### Murine KR158 SCCs form GBM-like tumor with heightened stemness and aggressive phenotype

The positive tumorigenicity of KR158 SCCs was evident through their intracranial implantation immediately after FACS isolation, resulting in the formation of brain tumors displaying hallmark histological features of GBM. These characteristics included infiltration and subpial spreading (***Fig. 6A-B***). Additionally, high levels of mitotic activity were evident, illustrated by cells in metaphase, anaphase, and telophase (***Fig. 6C, Fig. 6c1-3***). Furthermore, the presence of multipolar atypical mitotic figures was noted (***Fig. 6C, Fig. 6c4-6***). Additionally, the presence of red blood cells and perivascular aggregation was demonstrated (***Fig. 6C, Fig. 4c7-9***), along with well-defined pseudopalisading necrosis and presence of pyknotic nuclei (***Fig. 6D***). While tumors derived from FCCs also exhibited these classic morphological features (***Fig. 6E-F***), the pattern of infiltration seems less dispersed. The differentiation between these tumors was further illustrated by bulk RNA sequencing, revealing a distinct clustering pattern through PCA (***Fig. 6G***). Notably, the FCC cluster appeared more compact in comparison to the more dispersed SCC fraction, consistent with the observations from scRNA sequencing (***Fig. 2A***). This pattern implies a higher degree of cellular diversity within the SCC lineage compared to the FCC lineage. GSEA indicated an enrichment of stemness genes in SCC-derived tumors (***Fig. 6H, Table 17***). The SCC tumors also displayed higher transcriptional expression of genes associated with cell migration, cell motility, and ECM receptor interaction (***Fig. 6I-K, Tables 18-20***). Also, the SCC derived tumors were found to be enriched with genes regulating lipid metabolism (***Fig. 6L, Table 21***). Crucially, from a functional perspective, these SCC-derived tumors were linked to a more aggressive phenotype. This was evidenced through longitudinal monitoring of tumor volumes, measured using an IVIS Spectrum imaging system that captures bioluminescence associated with luciferase activity. Tumors generated by the intracranial transplant of luciferase-expressing SCCs showed faster progression compared to the other groups (***Fig. 6M***), resulting in shorter survival time (***Fig. 4N)***.

**Figure 6.**
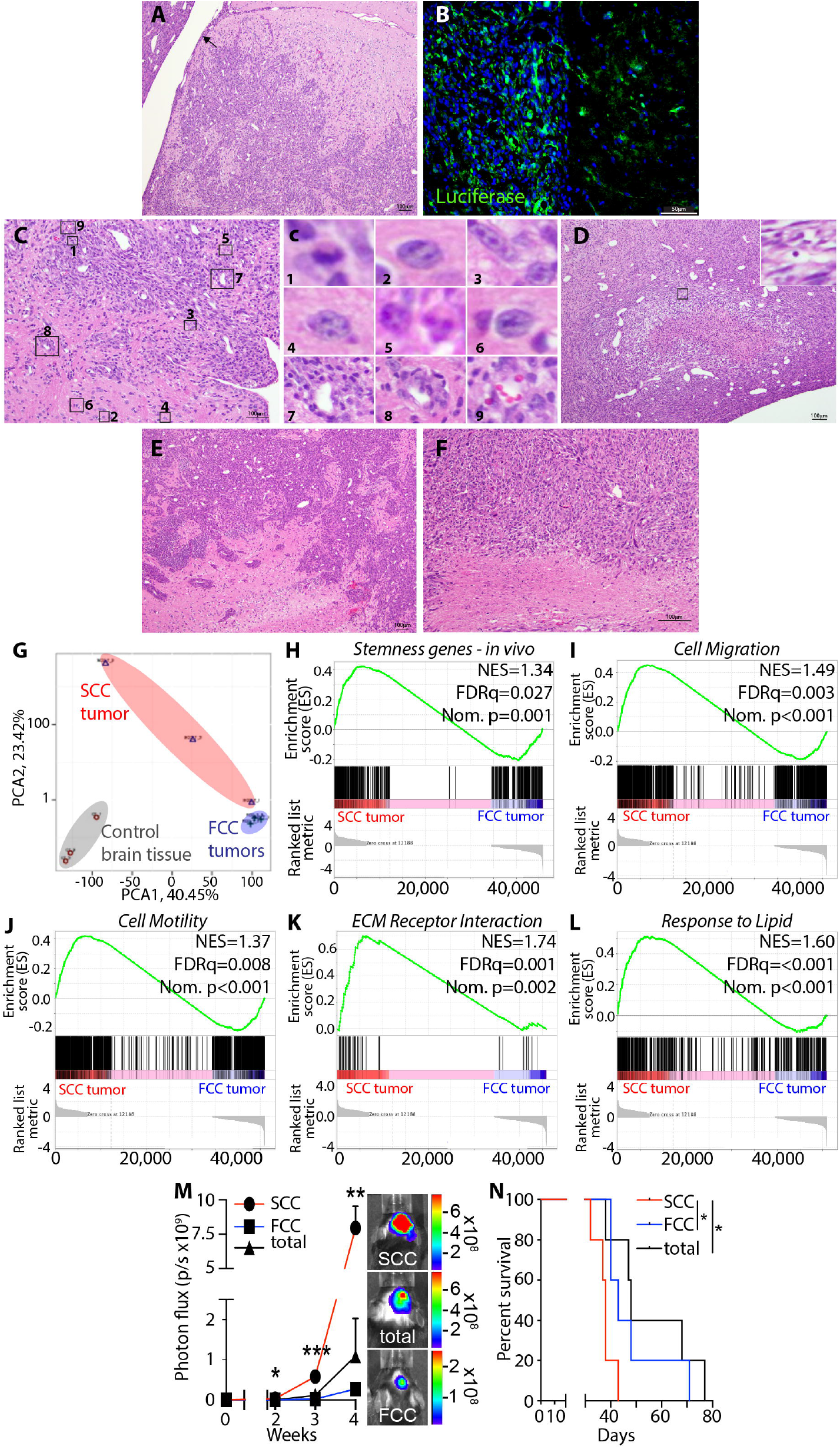
Murine glioma SCCs grown in the glioma sphere assay are tumorigenic with enhanced stemness. SCCs demonstrated ability to generate high-grade glioma like disease as seen by high level of tumor cell infiltration observed by H&E (**A**, → indicates subpial spreading) and luciferase labeling (**B**) as well as presence of mitotic figures (#1-6) and perivascular clustering (#7-9) (**C**). Panel (**c**) represents higher magnification insets of the panel C rectangles. (**D**) Pseudopalisading necrosis in SCC tumors (inset showing higher magnification of a pyknotic nucleus). H&E staining illustrated the ability of FCCs to also demonstrate classical high-grade glioma-like histological features of infiltration (**E**) and pseudopalisading necrosis (**F**). **G**) PCA score plot from RNA sequencing performed on tumors derived from SCCs and FCCs and from control brain tissue (n=3), showed that these different tumor cell populations maintain transcriptome diversity upon tumor progression. GSEA of *in vivo* RNA seq data sets between tumors derived from SCCs and FCCs (n=3 per group) **H)** for enrichment of stemness gene signatures using gene signature from Wong *et. al*. ^*34*^; GO terms were used to determine the enrichment of the following gene signatures : **I)** cell migration (GO:0016477) **J)** cell motility (GO:0048870); **K)** enrichment of gene signatures corresponding to ECM receptor interaction were obtained using the KEGG database (KEGG mmu04512) and **L**) response to lipids (GO:0071396); FDR, false discovery rate; NES, normalized enrichment score; Nom., nominal. **M**) Intracranial tumor growth monitored using bioluminescence *in vivo* imaging capturing luciferase activity in immunocompetent mice implanted with SCCs and FCCs (n=5). (**N**) Kaplan-Meier survival curves of immunocompetent animals implanted with SCCs compared to FCCs or total unsorted cells. *p<0.05, log-rank test.

### Murine KR158 SCCs are more tolerant to TMZ compared to FCCs

Based on the aforementioned phenotype of SCCs and the tumors they generate, we hypothesized that these cells may be less sensitive to chemotherapy. To evaluate the impact of the chemotherapeutic agent TMZ on the viability of freshly sorted SCCs, FCCs, and total unsorted tumor cells, we performed an *in vitro* PI incorporation assay. Similar to human GBM, murine SCCs displayed lower sensitivity to TMZ than the other cell populations (***Fig. 5A-B***). The fluorescent-based CyQuant assay was also used to compare cell numbers in the TMZ-treated cultures, revealing that SCCs had enhanced viability compared to FCCs (***Fig. 5C***). Additionally, the sphere forming frequency (SFF) assay indicated that SCCs exhibited significantly higher self-renewal activity in response to TMZ treatment than FCCs (***Fig. 5D-F***), further confirming their greater tolerance to the chemotherapeutic agent.

Together, these findings also support the conclusion that the KR158 cells adapted to the gliomasphere assay contain SCCs with properties and functions, including sphere forming ability, positive tumorigenicity, characteristic metabolic gene profile, and enhanced stemness and treatment resistance, similar to those described in human GBM ^16,19,20^. This study validates our immune competent murine model for investigating the relationship between tumor immune infiltrates and GBM cells, specifically SCCs.

## DISCUSSION

The intricate heterogeneity within GBM underscores its complexity and poses challenges in therapeutic approaches. The emergent understanding of CSCs, particularly slow-cycling cancer stem cells, supports their significant contribution to this heterogeneity and treatment resistance in GBM. Moreover, while the metabolic diversity within tumor cells that shapes their unique metabolic microenvironments has been identified ^1,19^, the precise influence of this heterogeneity on the immune landscape and disease progression remains largely elusive ^5^. This study is aimed to delineate and characterize GSCs, particularly SCCs, in an immunocompetent murine model of GBM. Our goal was to demonstrate that these cells, isolated from a specific murine model of GBM, mimic key phenotypic and functional traits observed in their human counterparts. Validating this paradigm in murine models will support their use in exploring the interactions between SCCs and the immune system, encompassing the study of their metabolic communications and dependencies. This deeper understanding of how tumor cell metabolic properties and diversity impact the tumor immune microenvironment will guide therapeutic strategies towards modulating this metabolic interplay.

GBM models played a crucial role in understanding tumor dynamics. Patient-derived models, patient-derived xenografts, murine models like syngeneic models, and genetically engineered mouse models (GEMMs) offer distinct advantages and limitations in investigating the role of GSCs within the TME, especially the tumor immune microenvironment. These diverse tools vary in their ability to mimic the human tumor microenvironment. Patient-derived *in vitro* models, including primary cell cultures and tumor organoids, offer relatively easy accessibility for experimental manipulations and high-throughput drug screening, molecular profiling, and mechanistic studies. However, these models may lack key aspects of the tumor heterogeneity and microenvironment complexity compared to patient-derived xenografts (PDXs), which maintain the molecular, histological, and architectural characteristics of the original tumor, thus closely mirroring the genetic landscape of the patient tumor and thus maintaining their clinical relevance. Though clinically relevant, PDXs require an immunocompromised system to prevent graft rejection, limiting the investigation of the role of the immune system in tumor progression and treatment response.

Conversely, syngeneic murine models can offer insights into tumor-stroma interactions, especially tumor-immune cell communication. However, recapitulation of the diffuse and infiltrative nature of GBM has been challenging to achieve in murine models. Therefore, presenting this unique property of GBM in mice is desirable to more accurately model tumor-stroma interactions and GBM cell behavior. Examples of murine models of GBM include GEMMs where tissue specific promoters are used for oncogenic transformation in neural stem and progenitor cells such as nestin-, GFAP-, CNPase-, and S100beta-positive cells ^61-69^. Other syngeneic models formed through various induction methods can also include GL261, CT2A, and KR158 ^70-79^. Together, these murine models provide a molecular insight into the impact of genetic mutations on disease initiation, progression, and treatment outcome. However, being more homogenetic in nature, they are less reflective of the intratumoral genomic and phenotypic heterogeneity of GBM TME ^80^. Another limitation is the type of architecture of the tumors they generate, with limited tumor infiltration of the brain parenchyma and more circumscribed edges, contrasting with the highly invasive architecture typically observed in GBM patients ^16,81^. Additionally, these models, notably GL261 and CT2A, exhibit marked immunogenicity with elevated MHC I expression, differing from the immunological profile observed in human pathology ^37,82-85^. In contrast, the KR158 model displays lower immunogenicity and demonstrates inherent resistance to checkpoint inhibition, mirroring characteristics found in human GBM ^38^. This particular trait could potentially render this model more relevant, especially in the evaluation of experimental immunotherapies. With the goal to overcome some of these limitations, especially in recapitulating tumor heterogeneity and the morphological characteristics of human GBM, including infiltration into the surrounding immune competent brain parenchyma, we adapted the KR158 cell murine model to the gliomasphere assay. This assay was developed to culture, enrich, and study GSCs, which form tumors recapitulating the key architectural characteristics of GBM, such as infiltration ^26,27,86,87^. This assay was designed to retain the tumor heterogeneity, including the presence of GSCs alongside their differentiated progenies, facilitating the study of tumor cell hierarchy and diversity, and performing functional assays, such as investigating self-renewal capacity, differentiation potential, drug response, and tumorigenicity. Gliomasphere cultures maintain the phenotypic and genetic characteristics of the original tumor ^4,27^ while also preserving stem cell-like properties. These conditions also conserve the self-renewal and differentiation abilities of GSCs, contributing to the diversity within the tumor.

Studies have shown that murine glioma cell lines, such as GL261 and CT-2A, when cultured as gliomaspheres (GL261-NS; CT-2A-NS), demonstrate an enrichment for cancer stem-like cells. This enrichment is notably greater compared to cells expanded under serum-containing monolayer adherent conditions. Histopathological examination of tumors generated from GL261-NS and CT-2A-NS revealed characteristics consistent with high-grade glioma, presenting an aggressive infiltrative phenotype within the murine brain parenchyma ^74,88^. Specifically, CT-2A-NS tumors manifest both distant and adjacent satellite lesions but lack the formation of pseudopalisading necrosis. Our study demonstrated that culturing KR158 murine glioma cells in the gliomasphere assay enriched for cells with greater stemness and ability to generate tumors exhibiting all the key morphological features of GBM, including a high level of infiltration and well-defined pseudopalisading necrotic areas. The malignant lesions reveal chromosomal heterogeneity, including the presence of atypical mitotic features especially multipolar mitotic figures. This phenomenon has been previously documented in GBM tissues and is known to significantly contribute to the induction of aneuploidy in glioblastoma cells. These findings further underscore the significance of the use of this model in understanding the disease ^89^.

Importantly, we also demonstrated the existence of SCCs in these cultures, which showed up-regulated stemness programs, positive tumorigenicity, and greater resistance to treatment, which are properties also defined in GBM patient-derived SCCs ^16,19,20^. Our previous investigations indicated in hGBM that SCC and FCC populations represent independent lineages with limited phenotypic and functional overlap, contributing to the heterogeneous landscape of GBM ^20^. The current study also supports such a hypothesis, as tumors originating from SCC or FCC and composed by their respective progenies demonstrate significantly different transcriptomes (***Fig. 6G***), treatment responses (***Fig. 7***), and disease projections (***Fig. 6M-N***). Although both cellular fractions displayed invasive characteristics, their infiltration patterns appeared distinct. The SCC group exhibited highly dispersed and individually penetrating cells, while the FCC tumors showed a more collective yet diffused invasion with formation of satellite lesions nearby into the TME. Of note, this observation is qualitative rather than quantitative, supporting the need for further investigation to elucidate the differences in diffusion patterns and the underlying mechanisms regulating these properties. Finally, the exact vertical and horizontal hierarchical relationship between SCCs and FCCs remains to be conclusively established, possibly necessitating experiments involving lineage tracing, followed by comprehensive functional and phenotypic analyses conducted side by side. Furthermore, the considerable gene expression dispersion among SCC tumors, contrasting with the clustered expression pattern observed in FCC tumors in the UMPA and PCA plot (***Fig. 2A, 6G***), could suggest increased heterogeneity and plasticity in the SCC lineage. This notion is reinforced by the enriched stemness signature evident in SCCs (***Fig. 5-6***).

**Figure 7.**
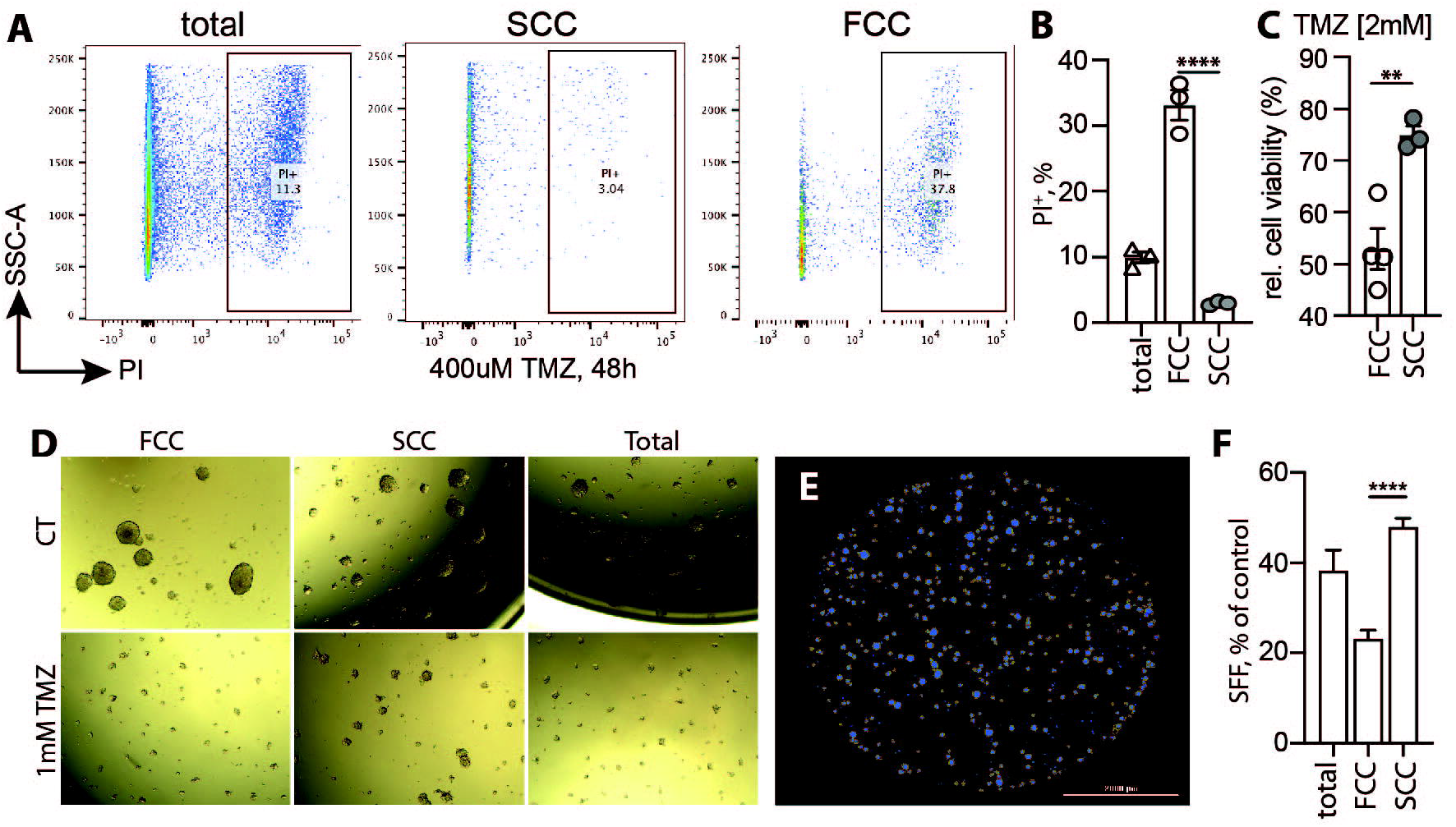
SCC KLuc cells are more resistant to TMZ with higher sphere forming frequency. **A-B)** PI incorporation assay within the tumor cells upon 48h of treatment with 400μM TMZ; Representative flow dot plots (**A**) with quantification of three independent experiments (**B**). **C**) Representative bar diagram of relative cell viability of SCCs and FCCs using the fluorescence-based CyQUANT Cell Proliferation assay upon exposure to 2mM TMZ, values represent mean +/-SEM. Brightfield (**D**) and fluorescent (DAPI) (**E**) images of gliomaspheres generated from the different KLuc cell populations treated with 1mM TMZ. **F)** Quantification of the data shown in panel D and E. The SFF was compared between the three cell populations (unsorted-total cells, SCCs and FCCs) treated with 1mM TMZ. Values represent mean +/-SEM, expressed as percentage of untreated conditions, n=19-20, ****p<0.001, one-way ANOVA with Tukey post-test).

## CONCLUSION

KR158 cells expanded in the gliomasphere assay in serum-free conditions replicate *in vivo* the diversity and heterogeneity of the tumor microenvironment in the context of a maintained immune contexture. Additionally, the infiltrative phenotype of the model combined with the existence of the slow-cycling cell paradigm further support its significance in investigating the involvement of GSCs and SCCs in GBM pathology and deciphering the complexity of the TME. Specifically, this model will provide an important platform to investigate the specific connections between SCCs and the immune system, and assess how these communications govern disease presentation, progression, and resistance to treatment. Importantly, this model presents an opportunity to evaluate strategies that target the interplay between SCCs and immune cells and assess their potential therapeutic effects.

## ABBREVIATIONS

(GBM): Glioblastoma
(CSCs): Cancer stem cells
(SCCs): Slow-cycling cells
(FCCs): fast-cycling cells
(GSCs): glioblastoma stem cells
(FACS): fluorescence-activated cell sorting
(IACUC): Institutional Animal Care and Use Committee
(OCT): Optimal Cutting Temperature compound
(TMZ): Temozolomide
(CTV): Cell Trace Violet
(PI): Propidium Iodide
(GEMMs): genetically engineered mouse models
(PDX): Patient derived Xenograft
(GSEA): Gene Enrichment Sequencing Analysis
(TME): Tumor Microenvironment
(PFA): Paraformaldehyde
(ANOVA): Analysis of variance formula
(KLuc): KR158-Luciferase

## AUTHOR CONTRIBUTIONS

Conceptualization: AC, CY, DAM, and LPD; Methodology: AC, CY, JLK, AS, DF, GT, MA and LPD; Validation: AC, CY, JH, JKH, MS, DAM, and LPD; Formal Analysis: AC, CY, JLK, JH, MS, and LPD; Investigation: AC, JLK, AS, DF, GT, MA, and LPD; Resources: DAM, and LPD; Data Curation: AC, JLK, CY and LPD; Writing –

Original Draft Preparation: AC and LPD; Writing – Review & Editing: JH, JKH, MS, and LPD; Supervision: LPD. All authors have read and agreed to this version of the manuscript.

## ACKNOWLEDGMENTS

This research was partially supported by the National Institute of Health (1R01NS121075, 1R21NS116578 to LPD).

## DATA AVAILABILITY STATEMENT

Data supporting the findings within this study are presented within the article and are available from the corresponding author upon reasonable request.

## SUPPLEMENTAL FILES

**Supplemental Video 1. Scratch-wound assay**. Time-lapse imaging shows the migration capacity of KR158 cells expanded in the gliomasphere assay during multiple passages prior to conducting the scratch assay. The video comprises 283 frames captured every 10 minutes and presented at a rate of 5 frames per second.

**Table 1.** Comparison of GSEA output: Stemness geneset ^34^ expression levels in tumors derived from KR158 cells cultured with serum versus GL261 cells.

**Table 2.** Comparison of GSEA output: Migration geneset (GO:0016477) expression levels in tumors derived from KR158 cells cultured with serum versus GL261 cells.

**Table 3.** Comparison of GSEA output: Cell motility geneset (GO:0048870) expression levels in tumors derived from KR158 cells cultured with serum versus GL261 cells.

**Table 4.** Comparison of GSEA output: ECM receptor interaction geneset (KEGG mmu04512) expression levels in tumors derived from KR158 cells cultured with serum versus GL261 cells.

**Table 5.** Comparison of GSEA output: SCC-related geneset ^19,20^ expression levels in tumors derived from KR158 cells cultured with serum versus GL261 cells.

**Table 6.** Comparison of GSEA output: Stemness geneset ^34^ expression levels in tumors derived from KR158 cells cultured in the gliomasphere assay versus KR158 cells expanded as monolayer in serum-containing conditions.

**Table 7.** Comparison of GSEA output: Stemness geneset ^35^ expression levels in tumors derived from KR158 cells cultured in the gliomasphere assay versus KR158 cells expanded as monolayer in serum-containing conditions.

**Table 8.** List of pathways up-regulated from the stemness gene network identified in KR158 cells cultured in the gliomasphere assay compared to adherent serum-containing conditions.

**Table 9.** Comparison of GSEA output (gliomasphere assay): Stemness geneset ^34^ expression levels in SCCs versus FCCs.

**Table 10.** Comparison of GSEA output (gliomasphere assay): Stemness geneset ^35^ expression levels in SCCs versus FCCs.

**Table 11.** Comparison of GSEA output (gliomasphere assay): SCC-related geneset ^19,20^ expression levels in SCCs versus FCCs.

**Table 12.** Comparison of GSEA output (gliomasphere assay): Migration geneset (GO:0016477) expression levels in SCCs versus FCCs.

**Table 13.** Comparison of GSEA output (gliomasphere assay): Cell motility geneset (GO:0048870) expression levels in SCCs versus FCCs.

**Table 14.** Comparison of GSEA output (gliomasphere assay): ECM receptor interaction geneset (KEGG mmu04512) expression levels in SCCs versus FCCs.

**Table 15.** Comparison of GSEA output (gliomasphere assay): Lipid catabolic process geneset (GO:0016042) expression levels in SCCs versus FCCs.

**Table 16.** Comparison of GSEA output (gliomasphere assay): Cellular response to lipid geneset (GO:0071396) expression levels in SCCs versus FCCs.

**Table 17.** Comparison of GSEA output: Stemness geneset ^34^ expression levels in tumors derived from SCCs versus tumors formed by FCCs, with cells expanded in the gliomasphere assay.

**Table 18.** Comparison of GSEA output: Migration geneset (GO:0016477) expression levels in tumors derived from SCCs versus tumors formed by FCCs, with cells expanded in the gliomasphere assay.

**Table 19.** Comparison of GSEA output: Cell motility geneset (GO:0048870) expression levels in tumors derived from SCCs versus tumors formed by FCCs, with cells expanded in the gliomasphere assay.

**Table 20.** Comparison of GSEA output: ECM receptor interaction geneset (KEGG mmu04512) expression levels in tumors derived from SCCs versus tumors formed by FCCs, with cells expanded in the gliomasphere assay.

**Table 21.** Comparison of GSEA output: Cellular response to lipid geneset (GO:0071396) expression levels in tumors derived from SCCs versus tumors formed by FCCs, with cells expanded in the gliomasphere assay.

